# Functionally convergent anti-predator morphologies arise through divergent cellular strategies in *Daphnia*

**DOI:** 10.64898/2026.08.04.742885

**Authors:** Shannon N. Snyder, Ethan B. Contreras, William A. Cresko

**Affiliations:** Institute of Ecology & Evolution, University of Oregon, Eugene, OR, 97403, USA; Department of Bioengineering, Knight Campus for Accelerating Scientific Impact, Eugene, OR, 97403, USA

**Keywords:** phenotypic plasticity, cell proliferation, inducible defenses, *Daphnia*, eco-evo-devo, predator-prey interactions, development

## Abstract

Phenotypic plasticity exemplifies how environmental signals can shape organismal development, yet the cellular mechanisms translating ecological cues into adaptive morphologies remain incompletely characterized. Many species of *Daphnia* (freshwater crustaceans) develop a diverse array of inducible defenses in response to predator chemical cues (kairomones), providing a tractable system for examining how ecological pressures are transduced through developmental mechanisms. *Daphnia lumholtzi* produce elongated head and tailspines in response to kairomones. How this occurs at the cellular level, through changes in cell size, proliferation, or both, is currently unknown. To address this question, we quantified the temporal dynamics of cell proliferation in *D. lumholtzi* across 72 hours following kairomone exposure using EdU incorporation (marking proliferating cells) and DAPI staining (to quantify total nuclei). Predator cue exposure induced a three-phase proliferative response: initiation within 24 hours (slightly increased proliferation and total cells), a transitional plateau at 48 hours (minimal effects in both treatments), and commitment by 72 hours (strong increases in both proliferation and cell accumulation). Headspines exhibited higher proliferation than tailspines, suggesting anterior-posterior developmental prioritization. Treated animals maintained smaller average cell sizes throughout the response, consistent with continuous addition of newly divided cells rather than cell enlargement. Unlike the delayed-division strategy in *D. longicephala* or bilayer formation in *D. pulex*, *D. lumholtzi* employs sustained hyperplasia, demonstrating that the same selective pressure produces similar ecological outcomes through mechanistically distinct developmental programs. Our findings bridge ecological signals with cellular responses, exemplifying eco-evo-devo integration whereby environmental pressures (predation) are transduced through developmental processes (cell proliferation dynamics) to generate adaptive and heritable morphological diversity across ecological and evolutionary timescales.

## Introduction

A central challenge in eco-evo-devo is understanding how environmental signals are transduced through developmental mechanisms such as cellular dynamics to produce adaptive phenotypes, connecting ecology, development, and evolution within a single causal framework (Brun Usan et al., 2025). Predator induced defenses have evolved as an adaptive form of phenotypic plasticity. Covariance of the phenotypic expression with the environment (Pigliucci, 2001) buffers prey from high-risk predator interactions (Auld et al., 2010; Batabyal, 2023; Ghalambor et al., 2007; Pettorelli et al., 2011; Via et al., 1995). The taxonomic breadth of predator-induced defenses is widespread (Tollrian & Harvell, 1990, 1999) and especially apparent in aquatic environments where organisms display remarkable convergent evolution of defensive responses (Agrawal et al., 1999; Riessen & Gilbert, 2019).

Although defense morphologies reduce predation success (Kruppert et al., 2019; Paplauskas et al., 2024), they are energetically expensive to produce, thereby creating trade-offs that affect fecundity and survivorship (Jin et al., 2024). As such, theory predicts that phenotypic plasticity should evolve as an advantageous strategy when predation pressure varies across spatial and temporal scales (Auld et al., 2010; DeWitt et al., 1998; Tollrian & Harvell, 1999), but can be lost in environments where predators no longer exert significant selective pressures. As such, induced predator defenses are a powerful system with which to determine *how* ecological signals are transduced through specific developmental pathways that influence evolutionary processes such as adaptation and diversification.

The freshwater crustacean, *Daphnia,* is a premier model organism for studying evolutionary and ecological dynamics, particularly of predator-induced defenses (Ebert, 2022). The *Daphnia* genus displays diverse defenses including helmets (*Daphnia cucullata*, Laforsch & Tollrian, 2004), crests (*Daphnia longicephala;* Horstmann et al., 2021), neckteeth (*Daphnia pulex*; Tollrian, 1993), and elongated spines (*Daphnia lumholtzi;*(Dzialowski et al., 2003). The morphological diversity across *Daphnia* species likely reflects differing predation pressures and highlights their evolutionary significance (Tariel et al., 2020). A number of studies have quantified phenotypic inheritance (Snyder et al., 2026), while others have characterized gene regulatory networks (Hales et al., 2017), but the cellular mechanisms that translate the environmental signal into a phenotypic outcome are less understood.

Defensive morphology development is dependent on tissue growth, which can be regulated by the division of cells (hyperplasia), the expansion of existing cells without division (hypertrophy), or a combination of both mechanisms (either sequentially or simultaneously deployed; Conlon & Raff, 1999; Neufeld & Edgar, 1998). Cell cycle checkpoints provide regulatory targets where environmental signals can modulate proliferation (hyperplasia) versus cell growth (hypertrophy; Crosby, 2008; Hartwell & Weinert, 1989). An alternative mechanism is endopolyploidy, where DNA replication occurs without cell division, increasing cell volume through the G1 phase (Edgar & Orr-Weaver, 2001; Lee et al., 2009). Endopolyploid cells are common in *Daphnia* epidermis (Beaton & Hebert, 1997), and are theorized to serve regulatory/secretory functions, directing neighboring diploid cells to undergo hyperplasia or hypertrophy by releasing neural molecules such as dopamine (Graeve et al., 2022).

Several distinct cellular and morphological strategies for defense formation have emerged across *Daphnia* species (Laforsch et al., 2004; L. Weiss et al., 2012). *Daphnia pulex* utilizes a bilayer formation strategy, in which the outer cellular layer secretes cuticle to form the neckteeth structure, supported by a hypertrophic pedestal formed by size increases of the inner layer. This strategy combines early hyperplasia (bilayer formation) with later cell cycle arrest (cell size increase of inner layer; Naraki et al., 2013). *D. longicephala* employs a delayed division strategy where predator-triggered cell cycle arrest allows increases in cell size before eventual division (Graeve et al., 2022), utilizing hypertrophy and delayed hyperplasia, markedly opposite the strategy implemented in *D. pulex.* In both species, large dopamine-filled, polyploid cells that likely release neurohormones are located near morphological defenses, potentially directing site-specific growth (L. C. Weiss et al., 2015). This indicates that although implementation differs, there is evidence of hyperplasia, hypertrophy and endopolyploidy within the *Daphnia* genus. If species-specific cellular mechanisms have evolved to achieve functionally similar outcomes, this reveals developmental lability, suggesting the decoupling of cellular mechanisms from ecological functions.

These findings raise a fundamental question: do different defense structures employ divergent cellular strategies across *Daphnia* species, or do conserved mechanisms produce quantitative variation? While both *D. pulex* neckteeth and *D. longicephala* crests employ temporal arrest of cell proliferation, *D. lumholtzi* elongates both head and tailspines to drive predator mishandling (Engel et al., 2014). The pointed spine geometry differs fundamentally from broad crests, suggesting spine elongation may favor active proliferation for linear tissue expansion. Additionally, whether both defensive spines on the same animal (head and tail) employ similar cellular strategies and temporal dynamics remains unexplored. A critical gap in our understanding is which cellular processes drive extensive morphological remodeling in response to environmental cues, and whether they are conserved across species.

To address this question, we focused on the cellular dynamics of induced head and tailspines in *D. lumholtzi* to compare our findings with previous work from other *Daphnia*. We quantified temporal dynamics (0-72h) of cell proliferation in predator-induced head and tailspines in order to characterize the cellular mechanism (active vs. delayed proliferation), compare anatomical spines (head versus tailspine), and assess conservation vs. divergence relative to other *Daphnia* species, thereby revealing developmental constraints on evolutionary diversification. We reveal how environmental signals are transduced through development to produce ecologically relevant outcomes and determine whether alternative cellular strategies can achieve similar ecological functions.

## Materials and Methods

### *Daphnia* source and laboratory maintenance

We used a *Daphnia lumholtzi* clone collected from Saguaro Lake, Arizona, USA in summer 2019 by Emily Williams (33.57470, −111.53690). From 2021 onward, animals were cultured in artificial lake medium (COMBO; Kilham et al., 1998) at 22°C under a 16:8 L:D cycle. We maintained 20-25 age-synchronized individuals per 250 mL beaker and fed them ad libitum with a 1:1 mixture of *Scenedesmus obliquus* and *Chlorella vulgaris*.

### Predator kairomone preparation

Predator-conditioned medium was prepared by culturing 10 threespine stickleback (*Gasterosteus aculeatus)* fish in 10 L COMBO for 3 days, feeding 20 adult *Daphnia* daily, then filtering (400 μm) and freezing. Control medium was prepared identically without fish. All preparation of kairomone was performed under an approved University of Oregon IACUC protocol.

### Treatment exposure timing

To ensure fine-scale age synchrony, we dissected neonates from the brood chamber at the last developmental stage, when animals are fully formed, and immediately placed individuals into either control or predator-conditioned medium until their respective endpoints (24, 48, or 72 hours) using randomized selection.

### Sample Collection Timeline and Rationale

We focused on the 0-72 hour post-dissection period because this represents a susceptible window for defense induction in our clone and parallels the examination period (0-72 hours after kairomone exposure during the sensitivity period) utilized in the Graeve et al. (2022) study on *D. longicephala*.

### Individual culture conditions

All animals were cultured individually in 30 mL of appropriate medium in 50 mL beakers at 22°C under a 16:8 L:D cycle. Animals were fed 5 mL of algae (85,000 cells/mL of *S. obliquus* and *C. vulgaris*, 1:1) daily. Attrition made large sample sizes difficult to achieve at the later timepoints. This could raise concerns about longer spines conferring a survival bias. However, our previous work (Snyder et al., 2026) demonstrated that there was not preferential survival based on spine length, arguing that attrition is unlikely to systematically bias the results. For the analysis, Bayesian hierarchical models are robust to unbalanced sample sizes, further justifying our analysis choice for main hypotheses.

### EdU and DAPI Incorporation Methodology

We used the Click-iT EdU Alexa Fluor 488 Imaging Kit (Thermo Fisher C10337) to label S-phase cells. EdU (20 μM) was added for 24-h pulses (0-24h, 24-48h, or 48-72h) to individual *Daphnia* in a 50 mL beaker with 30 mL of COMBO (Kilham et al., 1998). Animals were fixed in 4% formaldehyde/PBS (30 min), permeabilized (0.5% Triton X-100, 20 min), and processed according to manufacturer protocol. NucBlue Fixed Cell Stain ReadyProbes Reagent (Invitrogen R37606) was added prior to imaging at the manufacturer’s given ratio of 2 drops per mL.

### Cell Count Quantification

We quantified the number of cells and spine volume using Imaris imaging software (Bitplane AG, Oxford Instruments, Zurich, Switzerland). We defined two spines of interest: tailspine (from base of tail to spine tip) and headspine (from top of eye to spine tip). We validated automated cell counting through manual threshold-based approaches as detailed in the supplemental materials.

### Statistical Analysis

#### Model Framework and Justification

We used Bayesian hierarchical models to estimate treatment effects, accounting for unbalanced sample sizes, overdispersion, and repeated measures. We analyzed EdU and DAPI counts separately rather than as a ratio to distinguish proliferation changes from total cell accumulation and avoid statistical complications of ratio variables (spurious correlations, violated assumptions). Volume was log-transformed and included as a covariate. See supplemental materials for additional justification.

We employed a negative binomial error distribution rather than a Gaussian distribution due to overdispersion, and because model comparison strongly favored the negative binomial over Gaussian for modeling count data (ΔELPD = 71.8, SE = 34.4).

#### Priors for the Bayesian models

We used weakly informative priors, with the intercept prior set separately for each response to match its scale: Normal(5.5, 1) for the DAPI model and Normal(4, 1) for the EdU model, corresponding to approximately 245 and 55 cells on the response scale. Both models used Normal(0, 0.3) for fixed effects, Exponential(1) for random effect standard deviations, and Gamma(2, 0.1) for the negative binomial shape parameter. The treatment coefficient was given a Normal(0.15, 0.4) prior, weakly centered on a positive effect (approximately +16% on the response scale) to reflect the expectation from previous *Daphnia* work that predator cues increase cellular activity in developing defenses. This prior is diffuse relative to that expectation, spanning roughly a 50% decrease to a 160% increase within two standard deviations, and does not constrain the sign of the estimated effect. Treatment effect size magnitudes, ranging from approximately −2 to +62 cells depending on timepoint, spine, and measure, are presented in Supplemental Figure 1. See supplemental materials for additional explanation.

#### Model specification

We modeled cell counts using a negative binomial distribution with a log link function. The fixed-effects structure included treatment (control vs. predator-exposed as a factor variable), time (24 h, 48 h, 72 h as categorical variable), and spine (head vs. tail as a factor variable). We also included log-transformed spine volume to control for size differences, which were orders of magnitude larger than cell counts. An interaction between treatment and time was also included to capture potential temporal variation in treatment effects. A random intercept for individual was specified to account for repeated measures across head and tailspines within the same organism. The full model formula was specified as the following in R:

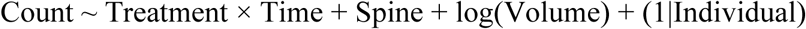

We tested Treatment × Volume and Treatment × Time × Spine interactions but excluded them due to overparameterization or lack of support in exploratory maximum-likelihood fits (p > 0.05).

#### Model fitting

We fit Bayesian models using the brms R package (version 2.22.0; (P. C. Bürkner, 2017)) with R version 4.5.0 (R Core Team, 2021). We used cmdstanr (version 0.9.0; (Gabry J & Johnson A, 2025)) as the Stan MCMC backend and ran 6 independent chains for 10,000 iterations each (4,000 warmup, 6,000 sampling).

#### Model Diagnostics

We assessed convergence using R^ statistics computed with the posterior package (Bürkner et al., 2025) and improved R^ methods (Vehtari et al., 2021). All parameters achieved R^ < 1.01, indicating chain convergence. We also evaluated effective sample sizes (ESS), which indicated sufficient independent samples for reliable posterior estimates (Bürkner et al., 2025). See Supplemental Table 1 for model diagnostics, and Supplemental Figure 2 for MCMC trace plots indicating convergence.

Visual inspection (PP-checks, bayesplot; Gabry et al., 2019) confirmed that models adequately captured the data, though we note slight underprediction in the EdU right tail (Supplemental Figure 3).

We explored alternative model structures and distributional families using Leave-One-Out Cross-Validation (LOO-CV) and Expected Log Pointwise Predictive Density (ELPD) using loo (Vehtari et al., 2017,Supplemental Table 2).

#### Effect Size Interpretation and Inference

We report P(Effect > 0), the posterior probability that an effect is positive, as the inferential statistic for all primary analyses. Where frequentist test statistics and p-values appear, they derive from exploratory maximum-likelihood fits used solely to screen candidate model structures, an approach we adopted to conserve computational resources. No primary inference in this study rests on those p-values. We categorize evidence as: P < 0.80 (inconclusive), P = 0.80–0.95 (directional), P > 0.95 (strong), as our own interpretive framework, not a universal standard. We visualized posterior distributions using bayesplot (Gabry et al., 2019).

We report 95% credible intervals (CrI), which can be interpreted as 95% probability that the true value falls within the interval given the data, incorporating all sources of uncertainty (sampling error, individual variation, model uncertainty). We report two effect size metrics: absolute effect size (mean difference in cell count, treated minus control), and relative effect size (percent change: [Difference/Control] × 100).

Model coefficients are presented in Supplemental Table 3.

#### Cell Size Analysis

To investigate whether cellular responses involved changes in average cell size, we calculated volume per cell (spine volume / total DAPI-positive cells) as an indirect proxy. Doing so assumes uniform cell density throughout the spine. Because volume per cell is a ratio of two measured quantities, it is disproportionately sensitive to spines in which few nuclei were segmented, which inflates the ratio. We therefore screened volume per cell using Tukey’s extreme-outlier rule, excluding values more than three interquartile ranges above the third quartile (an upper fence of 6,270 μm³). This removed six of 190 observations, all in the range 6,379–18,000 μm³, leaving n = 184. Fitting the same model to all 190 observations preserves the direction and rank order of every contrast but yields larger effects (for example, −28.9% rather than −18.2% at 72 hours in headspines), so the screened estimates we report are the more conservative of the two. We analyzed log-transformed volume per cell using a Bayesian hierarchical model:

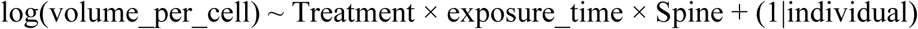

We used a Gaussian identity link family with a random effect to account for repeated measures. We specified the following weakly-informative priors: Intercept: Normal(8, 2) on log scale (corresponding to ∼3,000 μm³ on response scale), fixed effects: Normal(0, 0.5), random effect SD: Exponential(1), residual SD: Exponential(1). We fit models using brms (version 2.22.0) with 6 chains of 10,000 iterations (4,000 warmup).

#### Spine Volume Effects on Cell Counts

Because we were interested in temporal patterns where total cell counts and proliferation rates might move in opposite directions, we utilized spine volume as a covariate to account for size differences not dependent on treatment. As expected, cell counts scaled with volume due to larger tissue mass in larger anatomical structures. However, it is critical to separate treatment effects on cell density from effects mediated by size changes. We therefore analyzed spine volume as a predictor of cell counts and examined the treatment effect on volume.

We tested for Treatment × Volume interactions to verify that treatment effects were independent of body size. In exploratory maximum-likelihood fits, these interactions were non-significant for both DAPI (t = −1.39, p = 0.168) and EdU (t = −0.46, p = 0.648), justifying volume’s inclusion as an additive covariate rather than an interaction term.

#### Treatment Effect on Spine Volume

To test whether treatment affected spine volume independent of cellular changes, we fit a separate Bayesian model with log-transformed volume as the response variable:

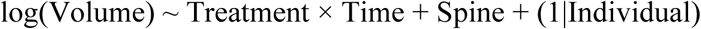

This model used a Gaussian distribution (appropriate for log-transformed continuous data), with priors: Normal(0, 2) for the intercept, Normal(0, 0.5) for fixed effects, and Exponential(1) for random effect and residual standard deviations. We ran 4 MCMC chains for 6,000 iterations (2,000 warmup, 4,000 sampling) with adapt_delta = 0.95. Model convergence and fit were assessed using the same diagnostics as cell count models (see above).

#### Reproducibility and Data Availability

All analyses were conducted in R version 4.5.0 (R Core Team, 2021) using brms version 2.22.0 (Bürkner, 2017) for Bayesian inference with the Stan backends cmdstanr (version 0.9.0; Gabry & Johnson, 2025) and rstan (Guo et al., 2015). Visualizations were created using ggplot2 (Wickham, 2016). Image analysis was performed using Imaris (Bitplane AG, Oxford Instruments, Zurich, Switzerland). Raw data and annotated code (R Markdown/Quarto documents) are available at github.com/shannonsnyder/daphnia-cell-proliferation.

## Results

### Predator-induced cell proliferation is weakly active at 24 hours, plateaus at 48 hours, and commits to rapid cell proliferation at 72 hours

We quantified both total cell accumulation (DAPI-positive nuclei) and proliferating cells (EdU-positive nuclei) across three developmental timepoints (24, 48, or 72 hours). We present un-modeled mean cell counts ± 1 standard error in Figure 1, illustrating temporal changes in proliferating cells and total cell counts as affected by treatment. Descriptive differences in mean cell counts and number of proliferating cells are apparent at 24 and 72 hours, with minimal differences at 48 hours (Table 1). The posterior distributions for both metrics are presented in Figure 2.

**Figure 1.**
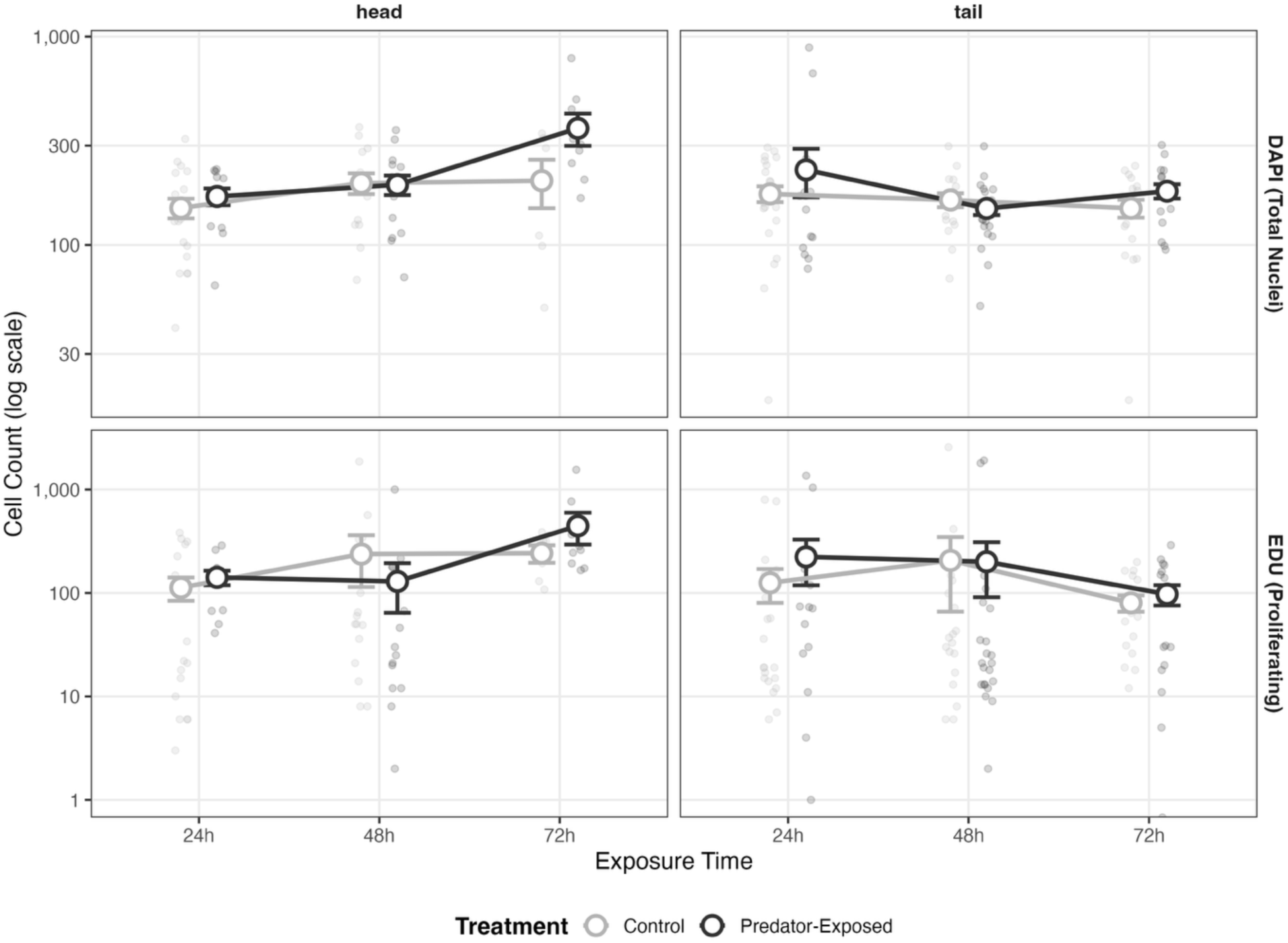
Temporal Dynamics of Cell Counts for Total (DAPI) and Proliferating (EdU) Cells. Longitudinal comparison of cell counts in head and tailspines of *Daphnia* following predator cue exposure over 72 hours. Top panels show DAPI-positive nuclei (total cells) and bottom panels show EdU-positive cells (proliferating cells in S-phase) on a log scale. Gray circles represent control animals; black circles represent predator-exposed animals. Large circles with error bars indicate group means ± standard error, while small translucent circles show individual observations. Note the log scale.

**Figure 2.**
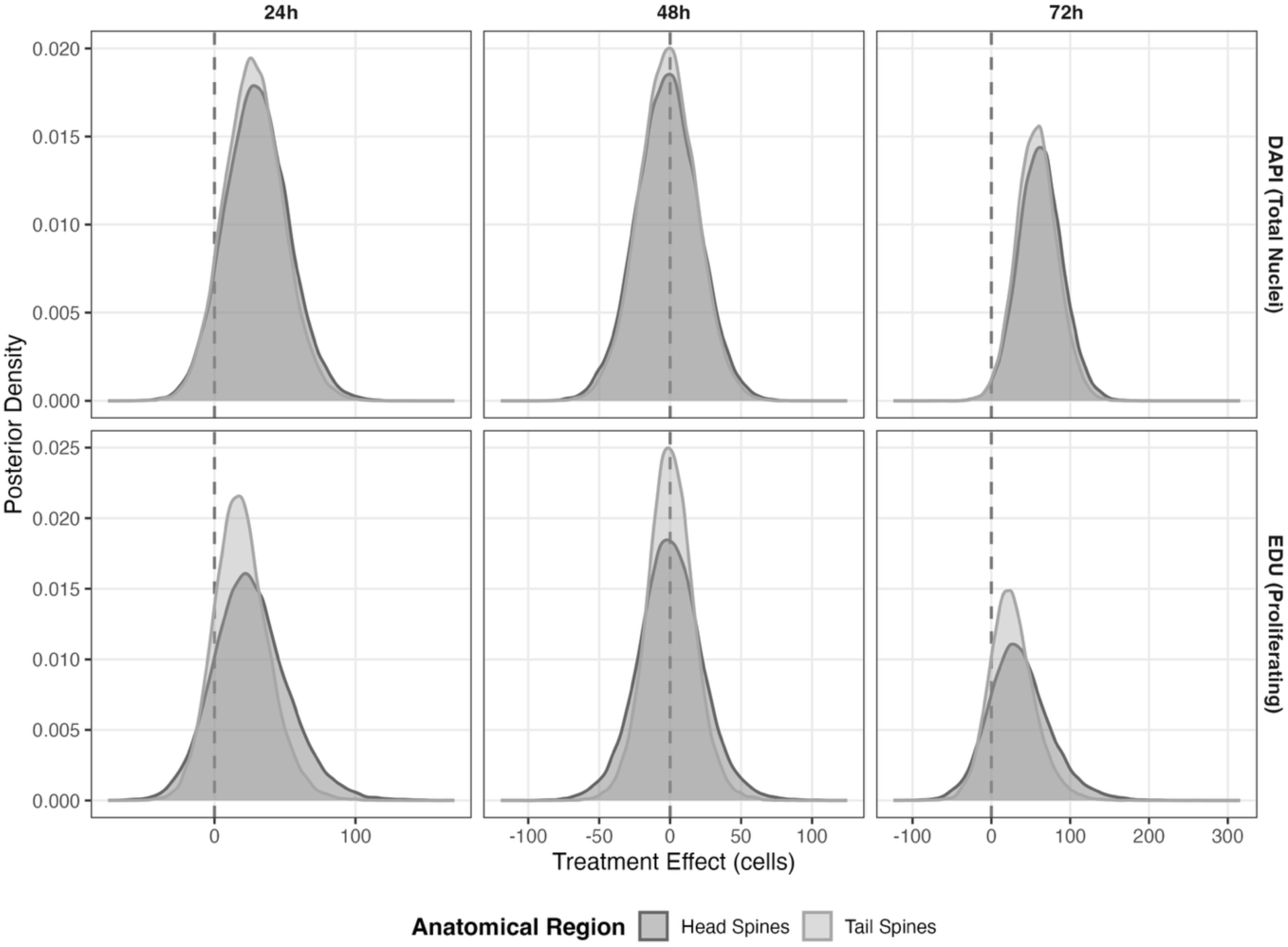
Posterior Distributions of Treatment Effects on Cell Counts. Bayesian posterior distributions showing the estimated treatment effects (predator-exposed minus control) on cell counts across three timepoints. Top row displays DAPI-positive cells (total nuclei); bottom row shows EdU-positive cells (proliferating cells). Dark gray distributions represent headspine; light gray represents tailspine. Vertical dashed lines indicate zero effect. The progression from 24h to 72h reveals a temporal pattern: initial trending effects at 24h (distributions spanning zero), transitional phase at 48h (distributions centered near zero), and strong positive effects at 72h (distributions shifted right of zero). By 72h the DAPI distributions sit clearly right of zero (P(Effect > 0) = 0.99), while the EdU distributions also shift right but with greater spread (P(Effect > 0) = 0.83), consistent with increased cell division during induced defense formation.

**Table 1.** Descriptive Statistics for Cell Counts and Volume by Treatment, Time, and Spine. DAPI-positive (total) and EdU-positive (proliferating) cell counts in head and tailspines of control and predator-treated D. lumholtzi at 24, 48, and 72 h post-exposure. N obs = number of observations; N ind = number of individuals. Values are mean, standard deviation (sd), standard error of the mean (se = sd/√N obs), and median. Volume in μm3.

| Treatment | Time | Spine | N obs | N ind | DAPI mean | DAPI sd | DAPI se | DAPI median | EdU mean | EdU sd | EdU se | EdU median | Volume mean |
| --- | --- | --- | --- | --- | --- | --- | --- | --- | --- | --- | --- | --- | --- |
| control | 24 | head | 20 | 18 | 150.50 | 72.82 | 16.28 | 137.0 | 112.55 | 127.41 | 28.49 | 63.5 | 305,565.0 |
| control | 24 | tail | 23 | 23 | 175.87 | 73.35 | 15.30 | 170.0 | 125.35 | 216.27 | 45.09 | 36.0 | 399,000.0 |
| control | 48 | head | 15 | 15 | 198.20 | 87.46 | 22.58 | 183.0 | 237.67 | 478.25 | 123.48 | 50.0 | 539,000.0 |
| control | 48 | tail | 18 | 18 | 164.44 | 53.83 | 12.69 | 163.0 | 206.33 | 595.71 | 140.41 | 35.0 | 382,277.8 |
| control | 72 | head | 6 | 6 | 203.50 | 130.64 | 53.33 | 201.5 | 242.33 | 114.22 | 46.63 | 255.5 | 567,833.3 |
| control | 72 | tail | 17 | 17 | 150.06 | 60.94 | 14.78 | 147.0 | 80.24 | 59.50 | 14.43 | 64.0 | 371,129.4 |
| treated | 24 | head | 12 | 12 | 171.00 | 54.41 | 15.71 | 179.0 | 141.33 | 78.33 | 22.61 | 141.5 | 310,416.7 |
| treated | 24 | tail | 15 | 15 | 229.53 | 231.91 | 59.88 | 172.0 | 223.33 | 405.93 | 104.81 | 73.0 | 249,353.3 |
| treated | 48 | head | 15 | 15 | 194.60 | 81.00 | 20.92 | 199.0 | 129.20 | 251.54 | 64.95 | 30.0 | 485,413.3 |
| treated | 48 | tail | 23 | 23 | 149.74 | 50.14 | 10.45 | 148.0 | 200.35 | 524.64 | 109.39 | 21.0 | 361,260.9 |
| treated | 72 | head | 9 | 9 | 363.22 | 191.78 | 63.93 | 304.0 | 445.22 | 454.46 | 151.49 | 260.0 | 678,333.3 |
| treated | 72 | tail | 17 | 17 | 181.29 | 58.91 | 14.29 | 178.0 | 97.24 | 89.05 | 21.60 | 85.0 | 390,764.7 |

Within 24 hours of predator cue exposure, treated animals showed directional increases in both total cell counts (Table 2) and proliferating cell counts (Table 3), with the raw trajectories shown in Figure 1 and the corresponding posterior distributions in Figure 2. In total cells (DAPI), headspines of unexposed animals averaged 168 cells versus 198 in predator-exposed animals, a 30-cell (+18%) increase (95% CrI [–12, 76], P(Effect > 0) = 0.91). Tailspines showed a similar pattern with 155 control cells versus 183 treated cells, a roughly 28-cell (+18%) increase (95% CrI [–12, 70], P(Effect > 0) = 0.91). Proliferating cells (EdU was added at the beginning of each sampling window and remained available for continuous incorporation; see methods) mirrored this pattern with proportionally larger effect sizes. Headspine proliferating cells increased from 97 (control) to 123 (treated), a 27% increase of 26 cells (95% CrI [–23, 83], P(Effect > 0) = 0.84). In tailspines, treated animals had 19 more proliferating cells (control 71 and treated 90), also representing a 27% increase (95% CrI [–17, 61], P(Effect > 0) = 0.84). While credible intervals cross zero at this timepoint, the 84–91% posterior probability favoring positive treatment effects, and the correlated increases across both cell types and both spines, suggest that predator-exposed animals are initiating cellular mechanisms contributing to defense development within the first 24 hours.

**Table 2.** Treatment Effects on Total Cell Count (DAPI) Reveal Temporal Variation in Total Number of Cells.

| Time | Spine | Mean Control | Mean Treated | Effect | P(+) | % Change |
| --- | --- | --- | --- | --- | --- | --- |
| 24h | head | 168.0 | 198.1 | 30.1 [-12.6, 76.2] | 0.91 | +17.9% |
| 24h | tail | 155.5 | 183.4 | 27.9 [-11.7, 70.2] | 0.91 | +17.9% |
| 48h | head | 172.5 | 170.5 | -2.0 [-45.2, 41.1] | 0.46 | -1.1% |
| 48h | tail | 159.6 | 157.8 | -1.8 [-42.1, 37.9] | 0.46 | -1.2% |
| 72h | head | 162.2 | 224.5 | 62.3 [8.4, 119.1] | 0.99 | +38.4% |
| 72h | tail | 150.1 | 207.7 | 57.7 [7.8, 110.3] | 0.99 | +38.4% |

Comparison of mean DAPI-positive cell counts between control and predator-treated *Daphnia* in head and tailspines across three timepoints. Effect values represent the difference between treated and control means (treated – control) with 95% credible intervals in brackets. P(+) indicates the probability that the treatment effect is positive. Percent change shows the relative increase in cell number following predator cue exposure. Notable increases occur at 24h (+17.9% in both spines) and 72h (+38.4% in both spines), with minimal change at 48h (-1.1% to -1.2%). The strong treatment effect at 72 hours (P(+) = 0.99) indicates a strong increase in the total number of cells in response to predator cues, consistent with defensive structure formation.

**Table 3.** Treatment Effects on Proliferating Cells (EdU) Indicates Active Proliferation.

| Time | Spine | Mean Control | Mean Treated | Effect | P(+) | % Change |
| --- | --- | --- | --- | --- | --- | --- |
| 24h | head | 96.5 | 122.3 | 25.8 [-22.6, 82.9] | 0.84 | +26.8% |
| 24h | tail | 71.1 | 90.2 | 19.1 [-16.6, 60.8] | 0.84 | +26.8% |
| 48h | head | 81.4 | 81.6 | 0.2 [-44.6, 46.7] | 0.50 | +0.2% |
| 48h | tail | 60.0 | 60.1 | 0.1 [-32.9, 34.1] | 0.50 | +0.2% |
| 72h | head | 109.9 | 145.5 | 35.6 [-35.6, 122.4] | 0.83 | +32.4% |
| 72h | tail | 80.9 | 107.1 | 26.2 [-26.0, 89.6] | 0.83 | +32.5% |

Comparison of mean EdU-positive cell counts between control and predator-treated *Daphnia* in head and tailspines across three timepoints. EdU incorporation indicates cells actively undergoing DNA synthesis (S-phase). Effect values represent the difference between treated and control means (positive indicates higher counts in treated animals) with 95% credible intervals in brackets. P(+) indicates the probability that the treatment effect is positive. Percent change shows the relative increase in proliferating cells following predator cue exposure. Minimal proliferation differences observed at 48h (P(+) = 0.50), with directional increases at 24h (P(+) = 0.84) and 72h (+32.4% in head, +32.5% in tail; P(+) = 0.83), suggesting proliferative response to predator cues during defensive structure formation in congruence with total cell increases.

At 48 hours, control and treated animals showed equivalent cell counts for both metrics (Tables 2 and 3, Figure 2). Total cells (DAPI) in headspines were approximately 170 in both groups (control was 172 cells whereas treated was 171 cells, with 95% CrI [–45, 41] and P(Effect > 0) = 0.46). There were comparable values in tailspines (control 160 and treated 158, with 95% CrI [–42, 38] and P(Effect > 0) = 0.46). Proliferating cells showed near-identical counts. Both control and treated headspines averaged 82 cells (95% CrI [–45, 47]), and both tailspine groups averaged 60 cells (95% CrI [–33, 34]). Notably, treated animals experienced sharper declines in both total and proliferating cells from the 24-hour timepoint. Treated headspines lost approximately 28 total cells and 41 proliferating cells from 24 to 48 hours, while control animals showed modest changes of +5 and –15 cells, respectively. These opposing trajectories, declines in treated animals versus stability in controls, resulted in convergence of the two groups at 48 hours.

After 72 hours of continuous predator cue exposure, treated animals demonstrated strong cellular responses in both total cell counts (Table 2) and proliferating cell counts (Table 3), summarized in Figures 1 and 2. Total cell counts (DAPI) increased by 62 cells (+38%) in headspines (control 162 and treated 225, with 95% CrI [8, 119] and P(Effect > 0) = 0.99) and by 58 cells (+38%) in tailspines (control 150 and treated 208, with 95% CrI [8, 110] and P(Effect > 0) = 0.99). With credible intervals excluding zero, these results represent strong evidence of treatment effects on total cell number. Proliferating cells (EdU) showed corresponding increases, though with wider uncertainty. Treated headspines contained 36 more proliferating cells than controls (control 110 and treated 146), a 32% increase (95% CrI [–36, 122] and P(Effect > 0) = 0.83). Tailspines showed 26 additional proliferating cells in treated animals (control 81 and treated 107), also a 32% increase (95% CrI [–26, 90], P(Effect > 0) = 0.83). While EdU credible intervals marginally cross zero, the consistent direction of effects with DAPI, which definitively shows increased total cell counts, makes it unlikely that proliferation patterns would contradict this evidence.

### Headspines contain more total and proliferating cells than tailspines at every timepoint

Across all timepoints (Table 4, Figure 3), total cell number (DAPI) was approximately 8% higher in headspines (P(Head > Tail) = 0.93 with 95% CrI [−2.5%, +19.7%]), and proliferating cells (EdU) were approximately 35% higher (P(Head > Tail) = 0.99 with 95% CrI [+5.5%, +74.2%]). Headspines contained more EdU-positive cells than tailspines in all six treatment × timepoint combinations, and more DAPI-positive cells in five of six (medians, Table 1). We therefore find a consistent anterior bias in cell number rather than an alternation between spines over time. Treatment effects were strongest at 72 hours in both spines, with control headspines containing 162 total cells compared with 225 in treated animals, and tailspines increasing from 150 to 208.

**Figure 3:**
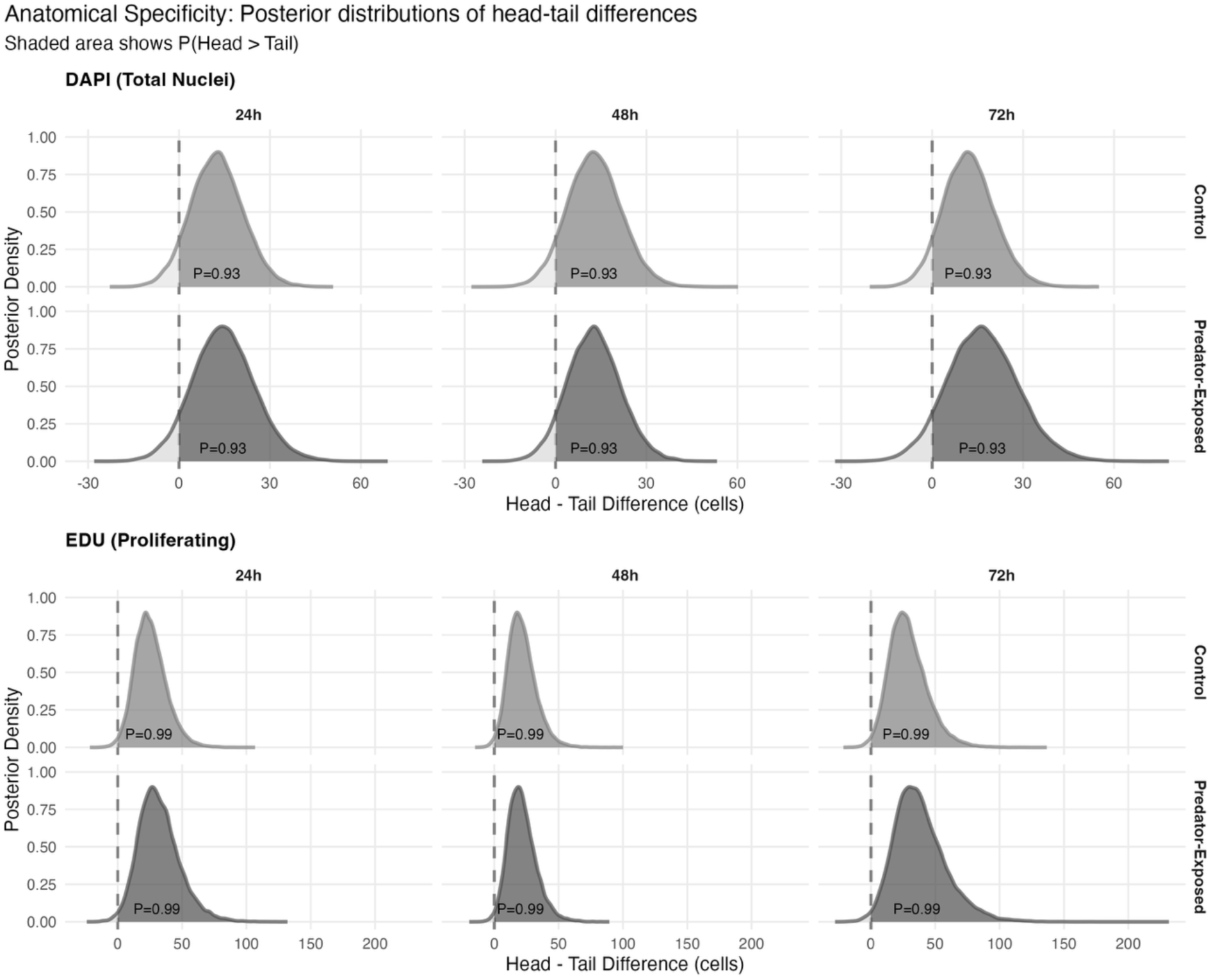
Posterior Distributions of Head and Tailspines Illustrate Preference for Headspine Elongation. Bayesian posterior distributions comparing cell counts between head and tailspines in control and predator- exposed *Daphnia*. Shaded area shows P(Head > Tail), representing the probability that headspines contain more cells than tailspines. Top panels show DAPI-positive nuclei (total cells); bottom panels show EdU-positive cells (proliferating cells). Vertical dashed lines indicate zero difference. For DAPI staining, headspines consistently show ∼10-20 more cells than tailspines across all conditions and timepoints (P = 0.93). For EdU staining, headspines contained 25–38 more proliferating cells in control animals and 32-85 more in predator-exposed animals (P = 0.99). The rightward shift and increased spread of EdU distributions in predator-exposed animals at 72h indicates preferential proliferation in headspines during defensive structure formation.

**Table 4.**
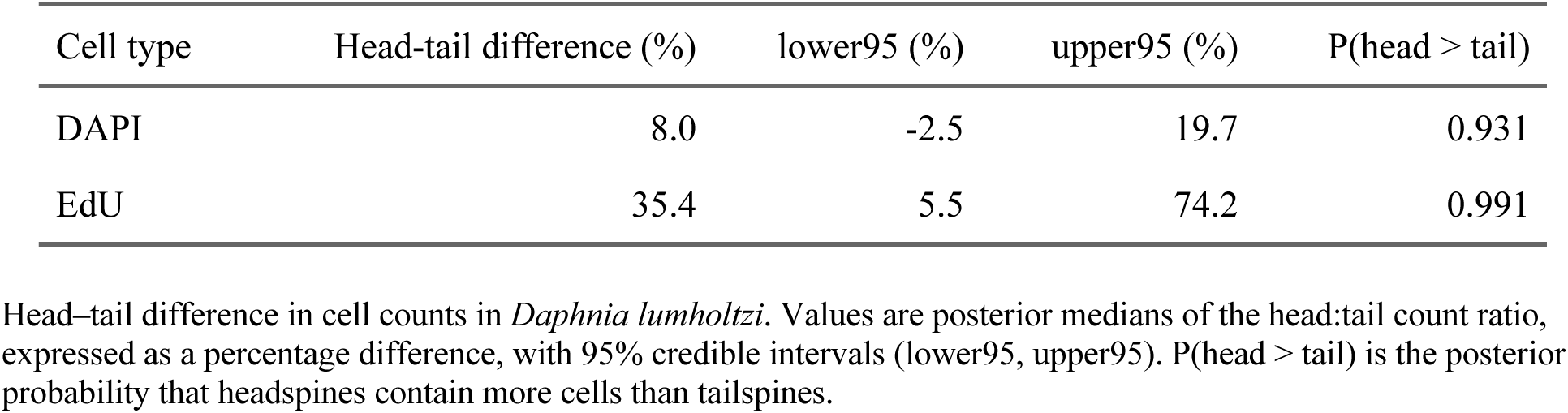
Headspines contain more total and proliferating cells than tailspines.

| Cell type | Head-tail difference (%) | lower95 (%) | upper95 (%) | P(head > tail) |
| --- | --- | --- | --- | --- |
| DAPI | 8.0 | -2.5 | 19.7 | 0.931 |
| EdU | 35.4 | 5.5 | 74.2 | 0.991 |

Head–tail difference in cell counts in *Daphnia lumholzi*. Values are posterior medians of the head:tail count ratio, expressed as a percentage difference, with 95% credible intervals (lower95, upper95). P(head > tail) is the posterior probability that headspines contain more cells than tailspines.

### Spine volume responds biphasically to predator cue exposure

For both measures, spine volume showed a positive relationship with cell number, with P(Effect > 0) = 0.99 for DAPI and P(Effect > 0) = 0.92 for EdU. This expected biological relationship of larger *Daphnia* containing more cells justified inclusion of volume as a covariate. The non-significant Treatment × Volume interaction (p > 0.16) indicates that treatment effects on cell counts are consistent across animal sizes, and that treatment does not differentially affect animals of different volumes. Treatment affected spine volume in a biphasic pattern, independent of cell counts (Table 5; Figure 4). At 24 hours, predator-exposed animals showed volume reduction (−48,276 μm³, −15.2%, P(Effect < 0) = 0.90). This effect weakened at 48 hours (−29,934 μm³, −7.0%, P(Effect < 0) = 0.70) and reversed to expansion at 72 hours (+37,881 μm³, +8.8%, P(Effect > 0) = 0.71), though all timepoints showed wide credible intervals crossing zero. These temporal volume changes, combined with the non-significant Treatment × Volume interaction, indicate that treatment affects cell proliferation directly, independent of size variation. Treatment does not differentially affect animals of different volumes, but does influence volume itself, with effects that change direction across the timepoints examined.

**Figure 4:**
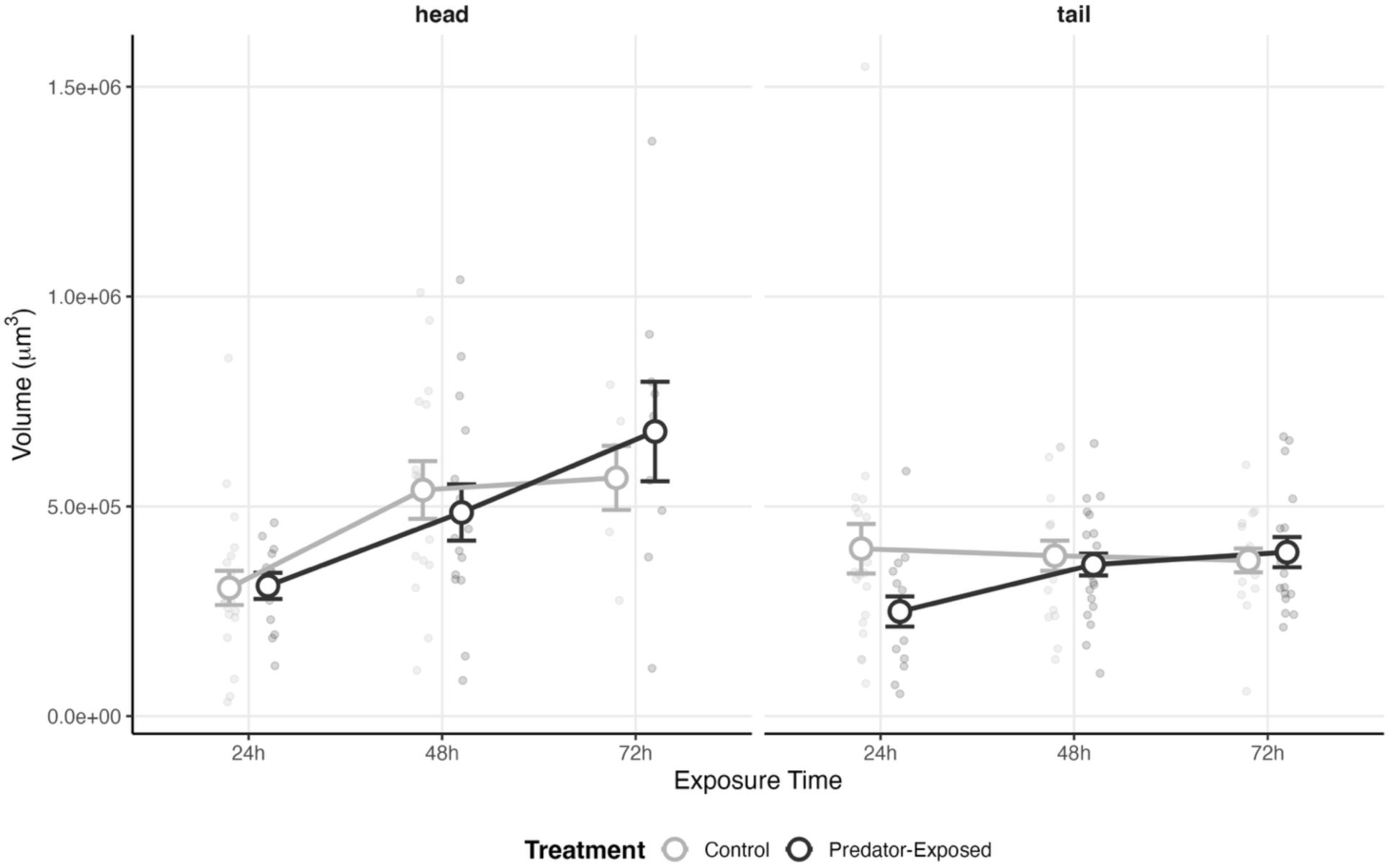
Treatment Effect on Spine Volume. Temporal changes in tissue volume (μm³) of head and tailspine in control and predator-exposed *Daphnia* over 72 hours. Gray circles represent control animals; black circles represent predator-exposed animals. Large circles with error bars indicate group means ± standard error, with individual observations shown as small translucent circles. Head and tailspines show broadly overlapping volume trajectories between control and treated groups; the modeled treatment effect is given in Table 5. Headspine volume increases from ∼305,000–310,000 μm³ at 24 h to ∼570,000– 680,000 μm³ at 72 h, while tailspine volume varies between ∼250,000 and ∼400,000 μm³ across timepoints.

**Table 5.** Treatment Effects on Spine Volume.

| Time | Mean control | Mean treated | Mean diff | lower_95 | upper_95 | Prob increase | Pct change |
| --- | --- | --- | --- | --- | --- | --- | --- |
| 24 | 317,807.2 | 269,530.8 | -48,276.46 | -121,922.63 | 26,339.67 | 0.10 | -15.19 |
| 48 | 429,262.6 | 399,329.1 | -29,933.55 | -140,808.76 | 78,018.24 | 0.30 | -6.97 |
| 72 | 429,272.7 | 467,153.9 | 37,881.19 | -99,212.63 | 175,635.90 | 0.71 | 8.82 |

Comparison of tissue volume measurements between control and predator-treated *Daphnia* over 72 hours. Mean diff represents the volume difference (treated minus control) in $\mu\text{m}^3$ , with 95% credible intervals (lower\_95, upper\_95). Prob increase indicates the posterior probability that treatment increases volume (therefore 0.10 represents 0.90 probability of decrease). Pct change shows the relative volume change following predator cue exposure (negative values indicate volume loss in treated animals; positive values indicate volume gain in treated animals).

### Predator-exposed animals maintain smaller average cell sizes

To investigate whether cellular responses involved changes in average cell size, we calculated volume per cell (spine volume / total DAPI-positive cells) as an indirect proxy, with modeled estimates reported in Table 6 and shown in Figure 5, and the underlying raw values in Supplemental Figure 4. At 24 hours, treated animals showed directional decreases in volume per cell. In headspines, control animals averaged 1,936 μm³ while treated averaged 1,685 μm³, a 13.0% reduction (P(decrease) = 0.81). This effect was more pronounced in tailspines. Treated animals (1,204 μm³) showed a 36.7% reduction compared to controls (1,904 μm³, with P(decrease) = 1.00 and CrI [−1233, −185]), providing strong evidence that predator cue exposure initially reduces average cell size, consistent with active proliferation adding smaller, newly divided cells to the population. At 48 hours, control and treated animals showed similar volume per cell in both spines (head 2,318 in control and 2,359 in treated, tail 2,319 in control and 2,362 in treated, with P(decrease) = 0.46 for both), indicating no clear directional effect during this transitional phase. By 72 hours, both spines trended toward smaller cells in treated animals. Headspines showed an 18.2% reduction (control 2,309 and treated 1,889 μm³, with P(decrease) = 0.80) and tailspines an 18.1% reduction (control 2,370 and treated 1,942 μm³, with P(decrease) = 0.85). While credible intervals cross zero at 72 hours, the 80–85% posterior probability favors smaller cells in treated animals.

**Figure 5.**
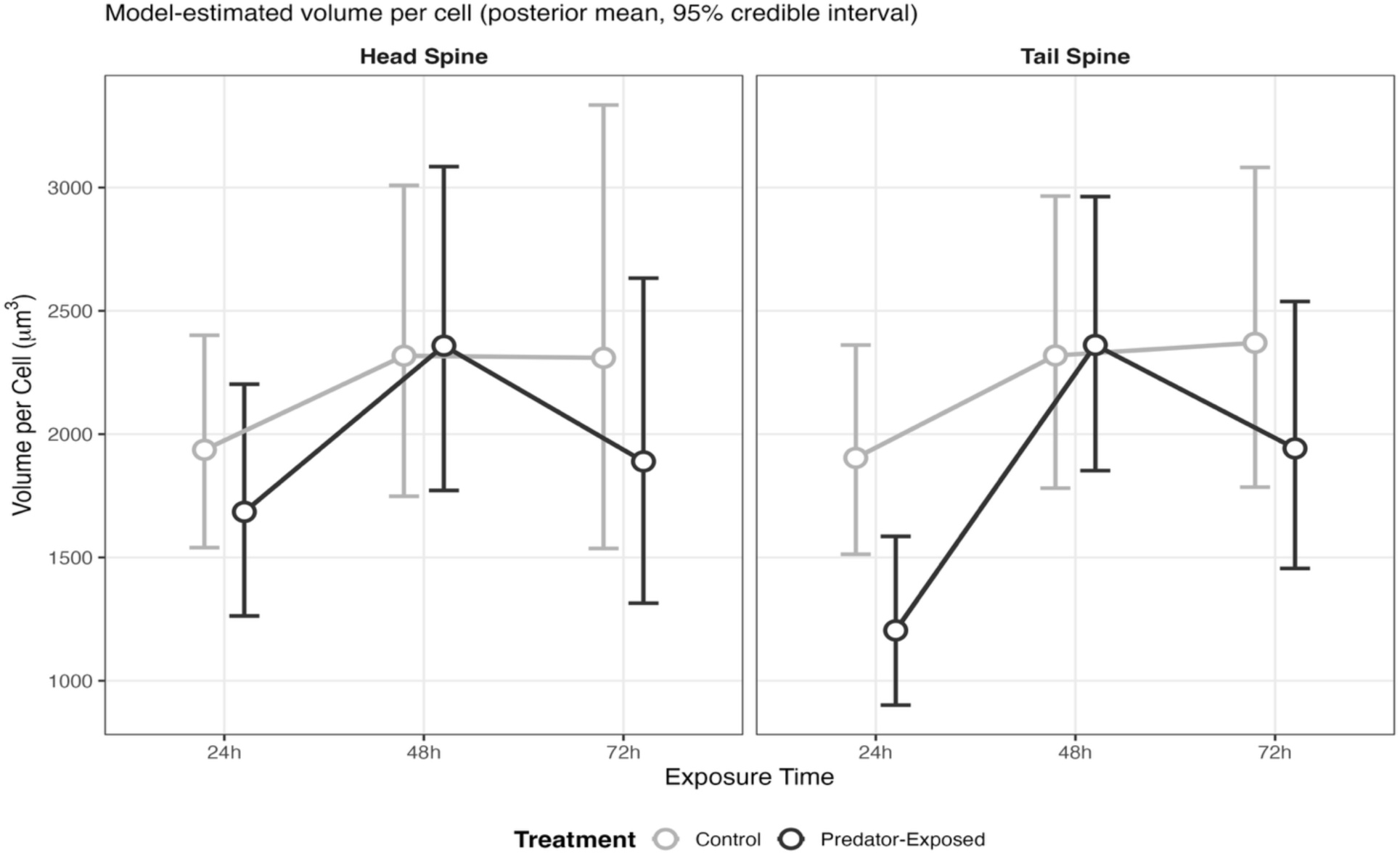
Treated Cell Volume Maintains Comparatively Smaller Cells Than Control Cells. Model-estimated volume per cell (spine volume divided by total DAPI-positive cells), an indirect proxy for average cell size. Points are posterior means of the fitted group mean and error bars are 95% credible intervals, from the Bayesian hierarchical model reported in Table 6. Control animals (light grey) showed higher volume per cell than predator-exposed animals (black) at the early (24 h) and late (72 h) timepoints, with the two groups converging at 48 h. Credible intervals overlap at every timepoint except the 24 h tailspine. The corresponding raw observations are shown in Supplemental Figure 4.

**Table 6.** Temporal Evidence of Treatment Induced Changes in Cell Volume.

| Time | Spine | Control | Treated | Difference | % Change | P(-) |
| --- | --- | --- | --- | --- | --- | --- |
| 24h | Head | 1936 ± 219 | 1685 ± 240 | -251 [-811, 334] | -13.0% | 0.81 |
| 24h | Tail | 1904 ± 216 | 1204 ± 174 | -699 [-1233, -185] | -36.7% | 1.00 |
| 48h | Head | 2318 ± 322 | 2359 ± 335 | 41 [-814, 910] | +1.8% | 0.46 |
| 48h | Tail | 2319 ± 304 | 2362 ± 283 | 43 [-765, 840] | +1.8% | 0.46 |
| 72h | Head | 2309 ± 460 | 1889 ± 337 | -420 [-1502, 549] | -18.2% | 0.80 |
| 72h | Tail | 2370 ± 329 | 1942 ± 277 | -428 [-1277, 389] | -18.1% | 0.85 |

Bayesian posterior estimates of treatment effects on volume per cell across timepoints and spines. Mean values are back-transformed from the log scale and are given as the posterior mean ± the posterior standard deviation of the fitted group mean. % Change is calculated from the two mean columns as (treated – control)/control, and agrees to within
0.5 percentage points with the posterior median of the per-draw ratio. Values in brackets are 95% credible intervals for the difference in $\mu\text{m}^3$ . $P(-)$ = posterior probability that treatment decreases cell volume; the 24 h tailspine value is 0.996, shown rounded.

## Discussion

Anti-predator morphologies are present across the *Daphnia* genus and could be governed by general cellular mechanisms facilitating tissue expansion, or by distinctive approaches tailored to the specific morphology. Here, we show that the sustained hyperplastic program documented in *D. lumholtzi* represents a mechanistically distinct strategy among characterized *Daphnia* species, in which environmental predator signals are transduced through active cell proliferation rather than through cell cycle arrest and hypertrophy. Fish cue exposure altered both the temporal dynamics and the anatomical allocation of cell proliferation, demonstrating that predator cues act as instructive signals that direct cellular programs within individuals.

### Temporal Dynamics Reveal Multi-Phase Developmental Programming

Within 24 hours of predator cue exposure, the cellular machinery supporting defensive spine development has been initiated, revealing a preference for cell division over strategies that suppress proliferation in favor of cell enlargement. Our 24-hour timepoint demonstrates consistent, though not decisive, coordination between cellular divisions and total cell accumulation, and by 72 hours strong positive effects in both metrics provide evidence that sustained predator cue exposure drives coordinated increases in cell division that expand cell populations.

Between 24 and 48 hours, minimal effects on both proliferation and total cell counts indicate a treatment-independent developmental plateau, which could be achieved mechanistically by stalling at G1 prior to commitment to cell division. We hypothesize that this plateau serves as a regulatory checkpoint that synchronizes developmental timing and assesses the temporal consistency of predator cues before committing resources to costly defensive structures, indicating that a singular exposure may not be sufficient for full defense formation. This generates the testable prediction that animals removed from predator medium at 48 hours should fail to execute the 72-hour proliferative response. Such tuning across developmental time would extend the fine modulation of cellular responses already demonstrated for predator cue concentration (Dennis et al., 2014) and exposure timing (Imai et al., 2009). Notably, predator-exposed animals show steeper decreases from 24 to 48 hours and larger increases from 48 to 72 hours than controls, so the comparable counts at 48 hours may mask an ongoing treatment effect. We also considered whether the plateau might represent a shift from active proliferation to cell growth, but treated animals maintained smaller cell sizes at 24 and 72 hours, while control cell size increased progressively. This pattern is consistent with the continuous addition of newly divided cells, supporting a hyperplasia-driven defensive strategy rather than hypertrophy.

Spine volume itself showed a biphasic response to predator cue exposure, independent of cell number changes. At 24 hours, treated animals exhibited volume reduction, suggesting initial resource reallocation toward cellular machinery rather than tissue expansion. This deficit attenuated by 48 hours and reversed to weak expansion at 72 hours. Because proliferation increased at both 24 and 72 hours while volume moved in opposite directions, these processes appear to operate through parallel pathways rather than volume mediating the cellular response. This supports a model where early predator detection triggers cellular priming before substantial tissue growth, with committed morphogenesis emerging only after sustained exposure.

### Compared to other *Daphnia*, *D. lumholtzi* has conserved processes and key differences

In our data, *D. lumholtzi* headspines consistently exhibited higher proliferation and total cell counts than tailspines, consistent with an anterior bias in the allocation of cell division. Both defensive structures respond to predator cue exposure, but developmental resources appear preferentially allocated to headspines. Spatially ordered proliferation has also been reported in *D. longicephala*, where crest formation begins in the ventral region and extends first in the cranial and then the dorsal direction rather than proceeding uniformly (Graeve et al., 2022). The large dopamine-containing cells implicated in defense morphogenesis are likewise anteriorly positioned, in the rostrum. Taken together, these observations (Graeve et al., 2022), and our work suggests that proliferation during defense formation is regionally patterned rather than uniform.

Common developmental mechanisms could explain this pattern. The morphogen gradient model posits that signaling molecules diffuse from a source, establishing concentration gradients that define proliferation zones (Gurdon & Bourillot, 2001; Wolpert, 1969). If such a morphogen exists in *Daphnia*, it could originate in the rostrum and drive the conserved anterior-to-posterior cascade, analogous to the Bicoid gradient in *Drosophila* (Struhl et al., 1989). Large dopamine-containing polyploid cells located in the rostrum of *D. longicephala* and *D. pulex* are hypothesized to release neurohormones regulating surrounding epidermis *(L. C. Weiss et al., 2015)*, so proximity to these cells could determine proliferation signal strength through local paracrine signaling. Dopamine also underlies life-history starvation responses (Issa et al., 2020), meaning that it could facilitate the integration of environmental signals before initiating costly defense formation. Testing whether dopamine signaling is conserved in *D. lumholtzi* spine development using immunological inhibition or dopamine receptor identification, is a priority for future work. Such a mechanism would integrate across biological scales, linking ecological predation risk to neural dopamine signaling, anterior-to-posterior proliferation gradients, and ultimately phenotypic response.

The active proliferation strategy we document in *D. lumholtzi* fundamentally differs from the delayed division mechanism reported in *D. longicephala* (Graeve et al., 2022), in which proliferating cells per unit area were reduced 48 hours after predator cue exposure, allowing cells to increase in volume before eventually dividing, a “grow first, divide later” strategy that maximizes cell size rather than cell number (Figure 6). Without direct cell size measurements, it also remains plausible that those cells are neither dividing nor growing but instead conserving energy for a later proliferative burst. Although we found no treatment-induced cellular arrest at 48 hours, both species experience reduced cellular activity at that timepoint, which could represent a conserved developmental strategy or limitation. *D. pulex*, which develops neckteeth in response to predator kairomones, demonstrates yet another pattern of initial cell proliferation followed by reduction (Naraki et al., 2013), though its temporal dynamics are less thoroughly characterized. Three species therefore deploy three distinct cellular strategies.

**Figure 6:**
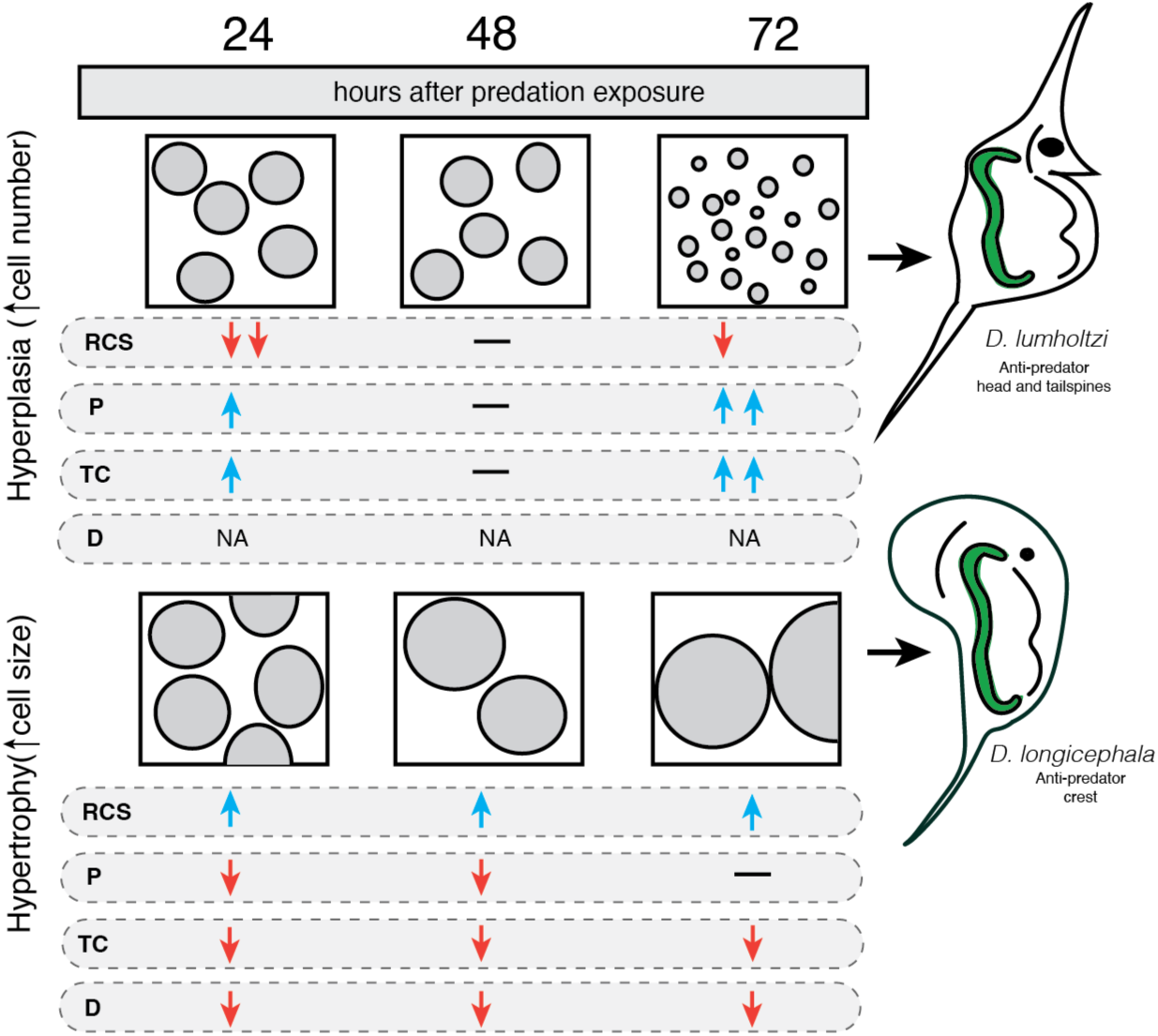
Summary Schematic. Species comparisons reveal distinct cellular strategies contributing to adaptive phenotypic plasticity. Summary schematic comparing developmental patterns (RCS = relative cell size (to control), P = proliferation, TC = total cell count, D = cell density) of *D. lumholtzi* (top: this study) and *D. longicephala* (bottom: Graeve et al., 2022) 24, 48, and 72 hours after the start of predation exposure. Blue arrows indicate increased values and red arrows decreased values in predator-exposed relative to unexposed *Daphnia*, with paired arrows marking the larger changes. Dashes in the RCS, P and TC rows indicate no detectable difference between treatments. Cell density (D) was not measured directly in our study, hence NAs.

We propose that these species-specific strategies are optimized for different morphological demands. Spine elongation in *D. lumholtzi* may be facilitated by rapid, active proliferation because hyperplasia along the growth axis provides building blocks for cell addition, producing a linear structure. In contrast, the broad crests of *D. longicephala* may benefit from larger cells contributing to an expansive surface. A potential driver of this divergence could relate to cuticle reinforcement documented in *D. pulex* and *D. longicephala* (Horstmann et al., 2021; Kruppert et al., 2016; Ritschar et al., 2020). Whether *D. lumholtzi* spines are reinforced remains unknown, but in our animals spine breakage is evident and not fatal, therefore spines may still mediate predator mishandling even if broken during the predator-prey interaction. Other species with crests and helmets likely cannot tolerate such structural failure. Thus, *D. lumholtzi* could prioritize proliferation at the expense of reinforcement, emphasizing length over thickness, a key deviation from broad helmet-like structures.

We found no evidence of delayed cell division; however, we cannot exclude early (0–24 hour) cellular arrest as in *D. longicephala*. Our earliest timepoint (24 hours) may have missed initial suppression at finer temporal resolution (2, 6, 12 hours). This mechanism would reconcile both approaches, showing that *D. lumholtzi* employs early delay followed by the late proliferation we observe.

### Implications for eco-evo-devo conceptualization

By characterizing another cellular pathway capable of achieving similar ecological functions, our findings are consistent with the hypothesis that the same selective pressures, such as predation, can produce functionally similar adaptive phenotypes through different cellular mechanisms (hyperplasia vs. hypertrophy) implemented across developmental timelines. A reaction-norm description would document that predator cues induce longer spines, plotting phenotype against environment without specifying how the change is produced, while mechanistic eco-evo-devo integration determines which cellular processes execute that response. Here we show that the same ecological challenge drives mechanistically distinct programs across closely related species. This decoupling of cellular mechanism from ecological function represents a fundamental principle of evo-devo. Development provides multiple routes to adaptive outcomes, with selection acting on phenotypes, while developmental mechanisms remain flexible for evolutionary innovation. Future studies could assess the generalizability of these principles across taxa.

## Data availability statement

The raw data and full analysis scripts that support the findings of this study are openly available on GitHub (https://github.com/shannonsnyder/daphnia-cell-proliferation).

## Author contributions

**Conceptualization and Methodology:** SNS and WAC conceived and designed the study. **Investigation:** SNS and EBC reared the animals and collected the imaging data. **Data Collection and Curation:** SNS curated the imaging measurements. **Formal Analysis:** SNS and WAC analyzed the data. **Visualization:** SNS created the data visualizations. **Writing – Original Draft:** SNS wrote the manuscript. **Writing – Review & Editing:** WAC provided revisions. All authors approved the final manuscript.

## Funding

This work was funded by the National Science Foundation Grant OPP-2015301 (WAC), University of Oregon Research Excellence funds (WAC), University of Oregon Office of the Vice President for Research and Innovation (OVPRI) seed funding (SNS and WAC). EBC received funding from the University of Oregon Graduate Students in Ecology and Evolution Student Organization and from the University of Oregon OVPRI.

## Conflict of Interest

The authors declare no conflict of interest.

## Acknowledgements

We would like to thank M. Lynch for his contribution of the *Daphnia lumholtzi* (SAG) utilized for this study. We would also like to thank Mark Currey for his assistance with stickleback husbandry and the Cresko lab for valuable feedback. We thank Emily Williams for collection and maintenance of *Daphnia* prior to their arrival in the Cresko lab. Finally, we graciously thank Adam Fries of the UO Genomics and Cell Characterization Facility for his vital help with fluorescent microscopy and cell quantification analysis.

## Notes

https://github.com/shannonsnyder/daphnia-cell-proliferation

### Competing Interest Statement

The authors have declared no competing interest.

