## Supplemental for "Functionally convergent anti-predator morphologies arise through divergent cellular strategies in *Daphnia*"

#### Supplemental Table 1. Model Diagnostics

| Model | Max_Rhat | Min_ESS_Bulk | Min_ESS_Tail | N_Divergent | Posterior draws |
|---|---|---|---|---|---|
| DAPI | 1.001462 | 5121 | 9793 | 0 | 36,000 |
| EdU | 1.000959 | 5254 | 10557 | 0 | 36,000 |

Convergence diagnostics for Bayesian models. Max_Rhat = maximum potential scale reduction factor ($<1.01$ indicates convergence). Min_ESS = minimum effective sample size for bulk/tail quantities. Models show good convergence (Rhat ≈ 1.00), adequate sampling (ESS > 400), and no divergent transitions (N_Divergent = 0). Posterior draws = total retained posterior samples (6 chains × 6,000 post-warmup iterations).

**Supplemental Table 2. Leave One Out Cross Validation**

| Model | ELPD | SE | P_LOO | LOOIC |
|---|---|---|---|---|
| DAPI | -1,085.334 | 17.55398 | 83.28456 | 2,170.668 |
| EdU | -1,077.794 | 20.53650 | 78.16152 | 2,155.589 |

Leave-one-out (LOO) cross-validation statistics for model predictive performance. ELPD (Expected Log Pointwise Predictive Density) indicates overall predictive accuracy, with higher (less negative) values representing better performance. SE shows the standard error of the ELPD estimate. P_LOO represents the effective number of parameters, indicating model complexity. LOOIC (LOO Information Criterion) provides a measure for model comparison, with lower values indicating better predictive performance.

**Supplemental Table 3. Model coefficients for DAPI and EdU Cell Counts**

| Model | Parameter | Estimate | Est.Error | Q2.5 | Q97.5 |
|---|---|---|---|---|---|
| DAPI | Intercept | 5.12006482 | 0.08882835 | 4.94537209 | 5.2945333 |
| DAPI | Treatmenttreated | 0.16294112 | 0.11906266 | -0.07134362 | 0.3963048 |
| DAPI | exposure_time48 | 0.02523503 | 0.11620919 | -0.20499270 | 0.2519557 |
| DAPI | exposure_time72 | -0.03749850 | 0.12369081 | -0.27934006 | 0.2057964 |
| DAPI | Spinetail | -0.07722398 | 0.05242263 | -0.18015720 | 0.0254664 |
| DAPI | log_volume_scaled | 0.13386072 | 0.04151151 | 0.05155785 | 0.2152974 |
| DAPI | Treatmenttreated:exposure_time48 | -0.17437512 | 0.15621559 | -0.47660727 | 0.1358990 |
| DAPI | Treatmenttreated:exposure_time72 | 0.16217834 | 0.16523498 | -0.15982318 | 0.4884955 |
| EdU | Intercept | 4.54755068 | 0.20994690 | 4.13556932 | 4.9586808 |
| EdU | Treatmenttreated | 0.23161186 | 0.22872764 | -0.22037807 | 0.6803693 |
| EdU | exposure_time48 | -0.17572397 | 0.21479049 | -0.59743786 | 0.2453611 |
| EdU | exposure_time72 | 0.12076455 | 0.22216792 | -0.31863621 | 0.5543630 |
| EdU | Spinetail | -0.30369650 | 0.12717879 | -0.55484643 | -0.0534250 |
| EdU | log_volume_scaled | 0.14995157 | 0.10531837 | -0.05667691 | 0.3584034 |
| EdU | Treatmenttreated:exposure_time48 | -0.23274664 | 0.24492212 | -0.71238151 | 0.2457684 |
| EdU | Treatmenttreated:exposure_time72 | 0.04211786 | 0.25074698 | -0.44628960 | 0.5316990 |

Posterior estimates from Bayesian hierarchical models for DAPI and EdU cell counts. Parameters show population-level effects on log-transformed counts. Estimate represents the posterior mean, Est.Error indicates the posterior standard deviation, and Q2.5/Q97.5 define the 95% credible intervals.

**Supplemental Figure 1. Treatment Effect Magnitudes**

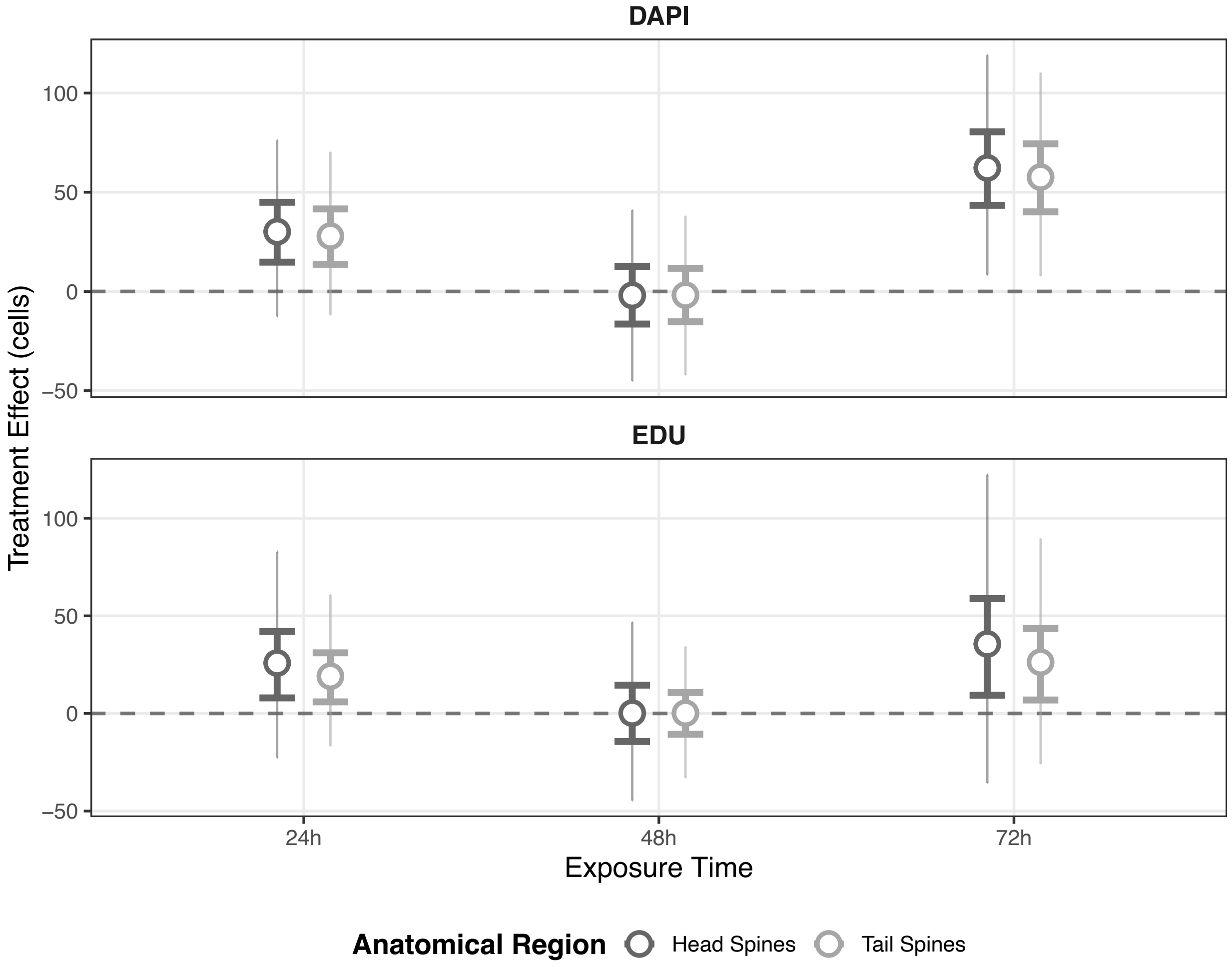


Temporal progression of treatment effects (predator-exposed minus control) on cell counts in head and tailspines. Top panel shows DAPI-positive cells (total nuclei); bottom panel shows EdU-positive cells (proliferating cells). Points represent posterior means with thick error bars showing 80% credible intervals and thin error bars showing 95% credible intervals. Dark gray circles indicate headspines, light gray circles indicate tailspines. Horizontal dashed line marks zero effect. Head and tailspines show remarkably similar response patterns.

### Supplemental Figure 2. MCMC Traces

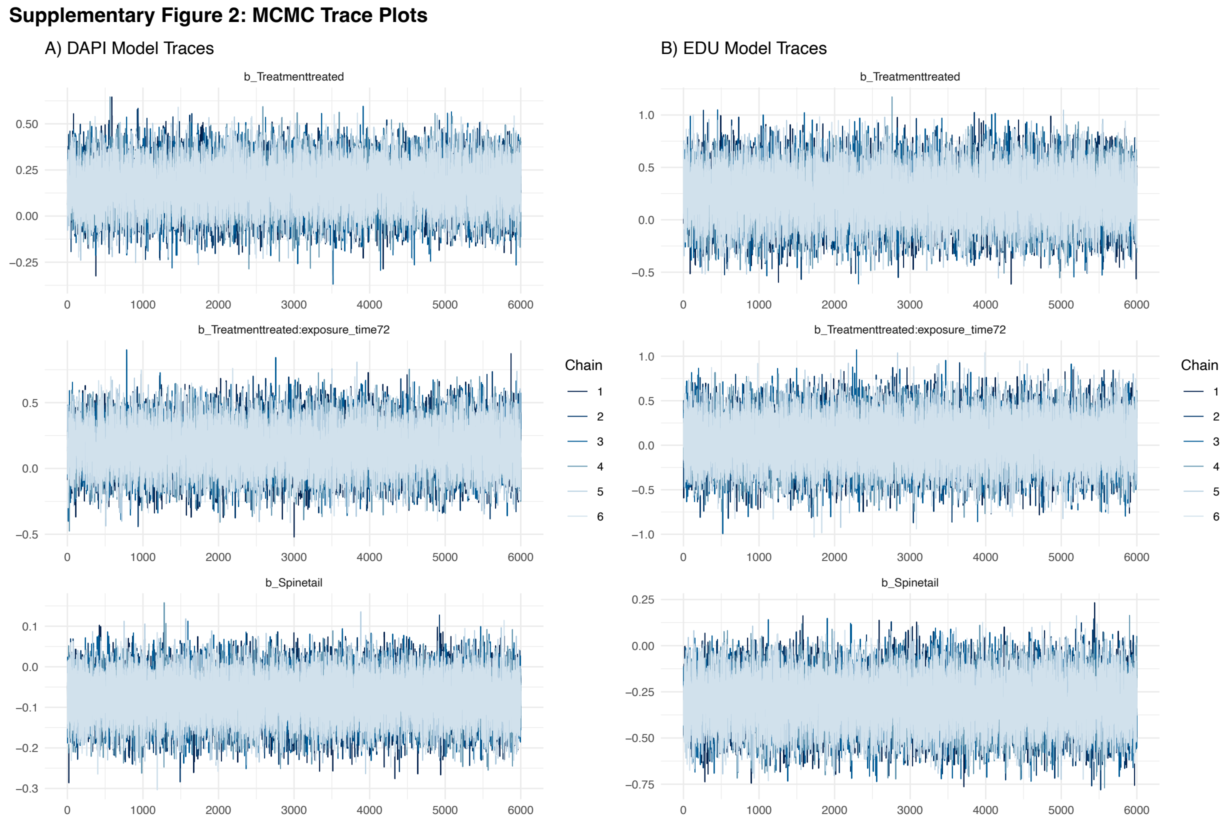


Markov Chain Monte Carlo (MCMC) sampling diagnostics for key model parameters in A) DAPI Model and B) EdU Model. Each panel shows trace plots for 6 independent chains (different shades of blue) across 6,000 iterations after warm-up. Parameters shown include: b_Treatmenttreated (main treatment effect), b_Treatmenttreated:exposure_time72 (treatment × time interaction at 72h), and b_Spinetail (spineeffect of tail vs. head). The overlapping, stationary chains exhibiting random walk behavior around stable means indicate good mixing and convergence. The wider variation in EdU model parameters (note different y-axis scales) reflects greater uncertainty due to higher biological variability in proliferation data compared to total cell counts.

**Supplemental Figure 3. Posterior Predictive Checks**

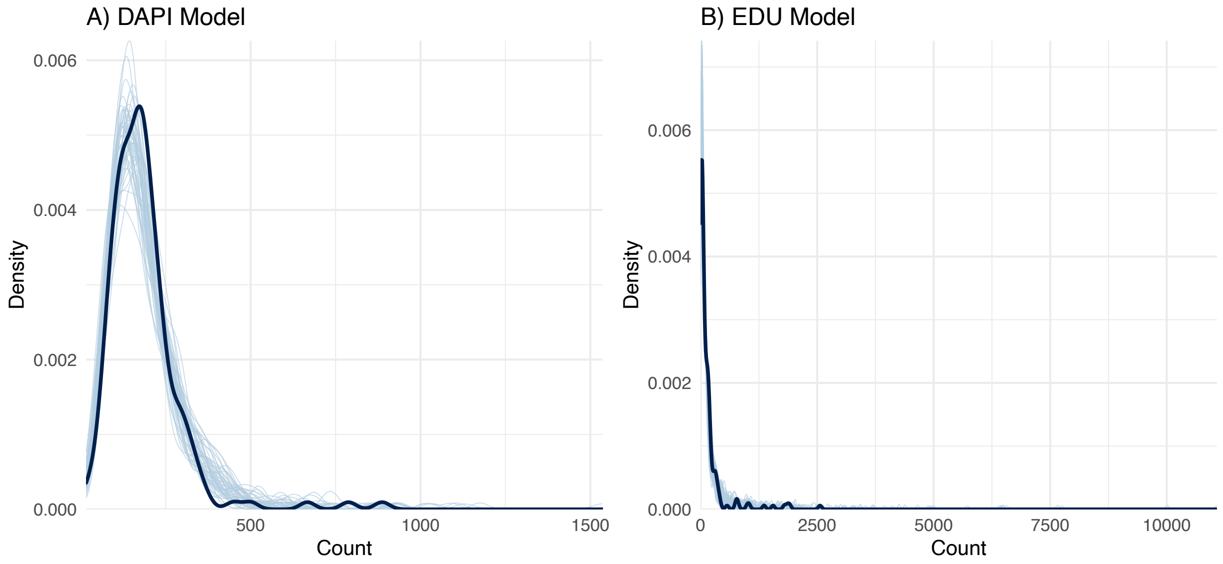


Model validation through posterior predictive distributions for A) DAPI Model and B) EdU Model. Dark blue lines show the density distribution of observed data; light blue lines represent 100 draws from the posterior predictive distribution. Close overlap between observed and predicted distributions indicates good model fit. The DAPI model (A) shows agreement between observed and predicted cell counts, with the model accurately capturing the peak around 150-200 cells and the right-skewed tail. The EdU model (B) similarly demonstrates good fit, accurately reproducing the sharp peak near zero (reflecting many samples with low proliferation) and the long tail extending to higher counts (representing samples with active proliferation). The wider distribution and longer tail in the EdU model reflects greater biological variability in proliferation rates compared to total cell counts. Both models successfully capture the overdispersion characteristic of count data, validating the choice of negative binomial distributions for both DAPI and EdU measurements.

**Supplemental Figure 4. Unmodeled Average Cell Volume**

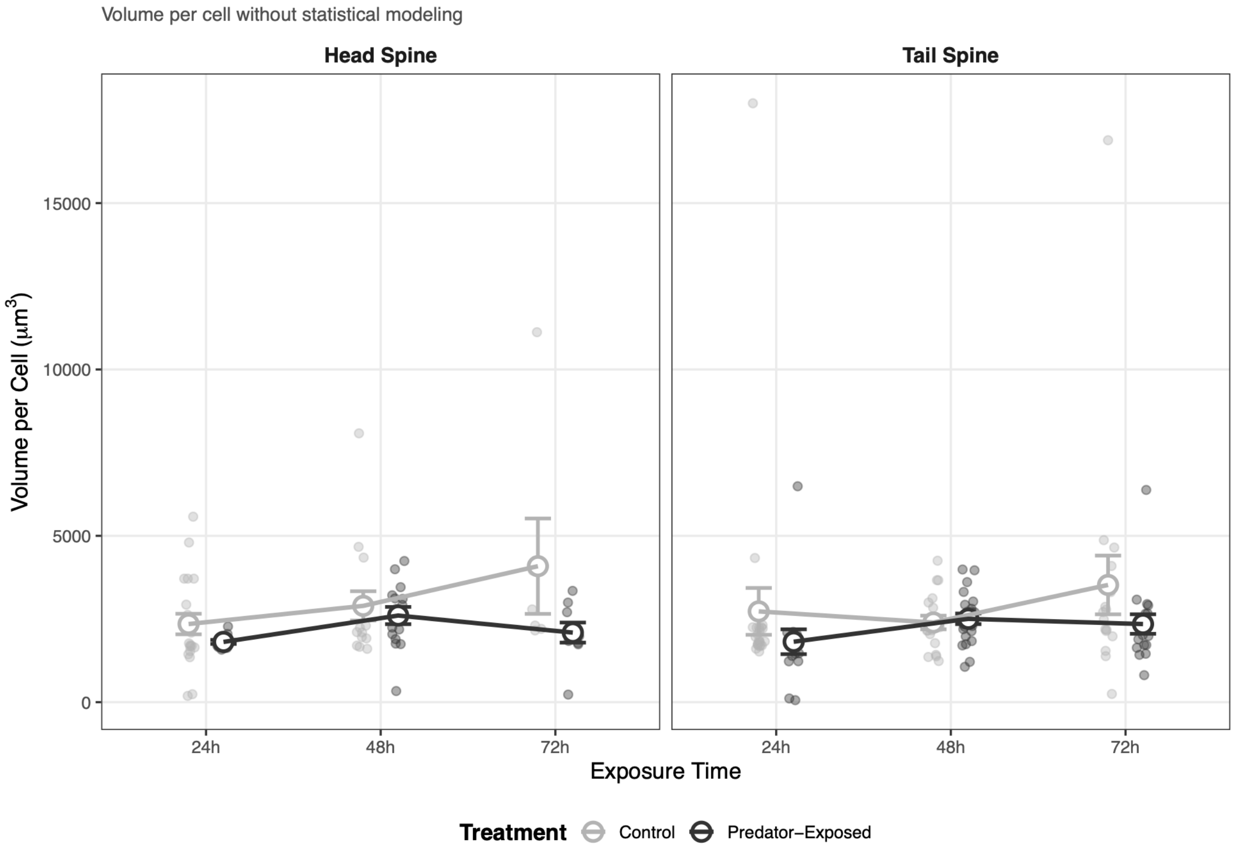


Volume per cell (spine volume divided by total DAPI-positive cells) serves as an indirect proxy for average cell size. Control animals (light grey circles) showed progressive increases in volume per cell across development, particularly in headspines from 48 to 72 hours. In contrast, predator-exposed animals (black circles) maintained relatively stable, smaller cell sizes throughout the 72-hour exposure period. This pattern is consistent with sustained active proliferation in treated animals, where continuous addition of small cells maintains lower average cell size, compared to a potential shift toward cell growth in controls. Points represent group means ± 1 standard error.
